# Negative Selection Allows DNA Mismatch Repair-Deficient Mouse Fibroblasts *In Vitro* to Tolerate High Levels of Somatic Mutations

**DOI:** 10.1101/2024.05.04.592535

**Authors:** Lei Zhang, Moonsook Lee, Xiaoxiao Hao, Joseph Ehlert, Zhongxuan Chi, Bo Jin, Alexander Y. Maslov, Albert-László Barabási, Jan H. J. Hoeijmakers, Winfried Edelmann, Jan Vijg, Xiao Dong

**Affiliations:** Institute on the Biology of Aging and Metabolism, University of Minnesota, Minneapolis, MN 55455, USA; Department of Biochemistry, Molecular Biology and Biophysics, University of Minnesota, Minneapolis, MN 55455, USA; Department of Genetics, Albert Einstein College of Medicine, Bronx, NY 10461, USA; Network Science Institute, Northeastern University, Boston, MA, USA; Department of Medicine, Brigham and Women’s Hospital, Harvard Medical School, Boston, MA, USA; Department of Network and Data Science, Central European University, Budapest, Hungary; Department of Molecular Genetics, Erasmus University Medical Center, Rotterdam, The Netherlands; University of Cologne, Faculty of Medicine, Cluster of Excellence for Aging Research, Institute for Genome Stability in Ageing and Disease, Cologne, Germany; Princess Maxima Center for Pediatric Oncology, Oncode Institute, Utrecht, The Netherlands; Department of Cell Biology, Albert Einstein College of Medicine, Bronx, NY 10461, USA; Center for Single-Cell Omics, School of Public Health, Shanghai Jiao Tong University School of Medicine, Shanghai, 200025, China; Department of Genetics, Cell Biology and Development, University of Minnesota, Minneapolis, MN 55455, USA; the Big Data Center of Sun Yat-sen Memorial Hospital, Sun Yat-sen University, Guangzhou, Guangdong 510123, China

**Keywords:** Aging, Mutation burden, Mutational signature, Single-cell whole-genome sequencing, DNA mismatch repair deficiency

## Abstract

Substantial numbers of somatic mutations have been found to accumulate with age in different human tissues. Clonal cellular amplification of some of these mutations can cause cancer and other diseases. However, it is as yet unclear if and to what extent an increased burden of random mutations can affect cellular function without clonal amplification. We tested this in cell culture, which avoids the limitation that an increased mutation burden *in vivo* typically leads to cancer. We performed single-cell whole-genome sequencing of primary fibroblasts from DNA mismatch repair (MMR) deficient *Msh2^-/-^* mice and littermate control animals after long-term passaging. Apart from analyzing somatic mutation burden we analyzed clonality, mutational signatures, and hotspots in the genome, characterizing the complete landscape of somatic mutagenesis in normal and MMR-deficient mouse primary fibroblasts during passaging. While growth rate of *Msh2^-/-^*fibroblasts was not significantly different from the controls, the number of *de novo* single-nucleotide variants (SNVs) increased linearly up until at least 30,000 SNVs per cell, with the frequency of small insertions and deletions (INDELs) plateauing in the *Msh2^-/-^* fibroblasts to about 10,000 INDELS per cell. We provide evidence for negative selection and large-scale mutation-driven population changes, including significant clonal expansion of preexisting mutations and widespread cell-strain-specific hotspots. Overall, our results provide evidence that increased somatic mutation burden drives significant cell evolutionary changes in a dynamic cell culture system without significant effects on growth. Since similar selection processes against mutations preventing organ and tissue dysfunction during aging are difficult to envision, these results suggest that increased somatic mutation burden can play a causal role in aging and diseases other than cancer.

## Introduction

Accumulation of somatic mutations has been proposed as a cause of aging and cancer since the 1950s (Failla, 1958; Szilard, 1959). DNA mutations occur spontaneously in every cell of an organism due to errors during repair or replication of a damaged DNA template (Vijg and Dong, 2020). However, apart from the very small fraction of mutations that are clonally amplified, the vast majority of mutations cannot be detected by bulk sequencing and require single-cell or single-molecule approaches. Using accurate single-cell whole-genome sequencing (scWGS) (Bohrson et al., 2019; Dong et al., 2017), somatic single-nucleotide variants (SNVs) have recently been found to accumulate with age in every human tissue or cell type analyzed, including lymphocytes (Zhang et al., 2019), hepatocytes (Brazhnik et al., 2020), epithelial cells (Huang et al., 2022), neurons (Lodato et al., 2018; Miller et al., 2022), and cardiomyocytes (Choudhury et al., 2022). Somatic SNV burden ranges from a few hundred to a few thousand mutations depending on cell type and age. While confirming the original hypotheses of somatic mutation accumulation with age, it remains unclear if an increased burden of somatic mutations, in the absence of clonal amplification, has functional consequences for cells and tissues at old age.

If mutation accumulation is indeed a cause of aging, one would expect an upper limit of mutations that cells can tolerate. Here we tested this using primary fibroblasts from a DNA mismatch repair (MMR)-deficient mouse model, i.e., *Msh2^-/-^* mice. The Msh2 (MutS homolog 2) gene encodes a protein that dimerizes with Msh6 and Msh3 proteins to make MutSα and MutSβ mismatch repair complexes, respectively, and is critical for correcting base mismatches and insertion or deletion mispairs during DNA replication (Li, 2008). Such mice are known to have highly increased somatic mutation frequencies and a greatly increased risk of cancer (de Wind et al., 1995; Hegan et al., 2006). The life span of a *Msh2^-/-^* mouse, 50% of which die within 6 months (Lin et al., 2004) is significantly less than that of a wild-type mouse in captivity, which typically lives to about 2-2.5 years, and the expression of *Msh2* is positively correlated with the maximum life span across different rodent species (Lu et al., 2022). The MMR deficiency would continually drive the generation of single-nucleotide variants (SNVs) and small insertions and deletions (INDELs) during passaging of these cells, allowing us to test a possible limit of tolerance *in vitro* (schematically depicted in **Fig. 1A**). The results show no such limit for SNVs up until at least ∼30,000 SNVs per cell, i.e. far exceeding the number of SNVs observed in most tissues upon normal aging. INDEL accumulation, however, reached a limit at <500 and ∼10,000 INDELs per cell in control and *Msh2^-/-^*cells respectively. Our results also indicate a strong negative selection against deleterious SNVs and INDELs, suggesting that somatic mutations can adversely affect cell function *in vivo* where selection for a fitness advantage is rarely possible.

**Figure 1.**
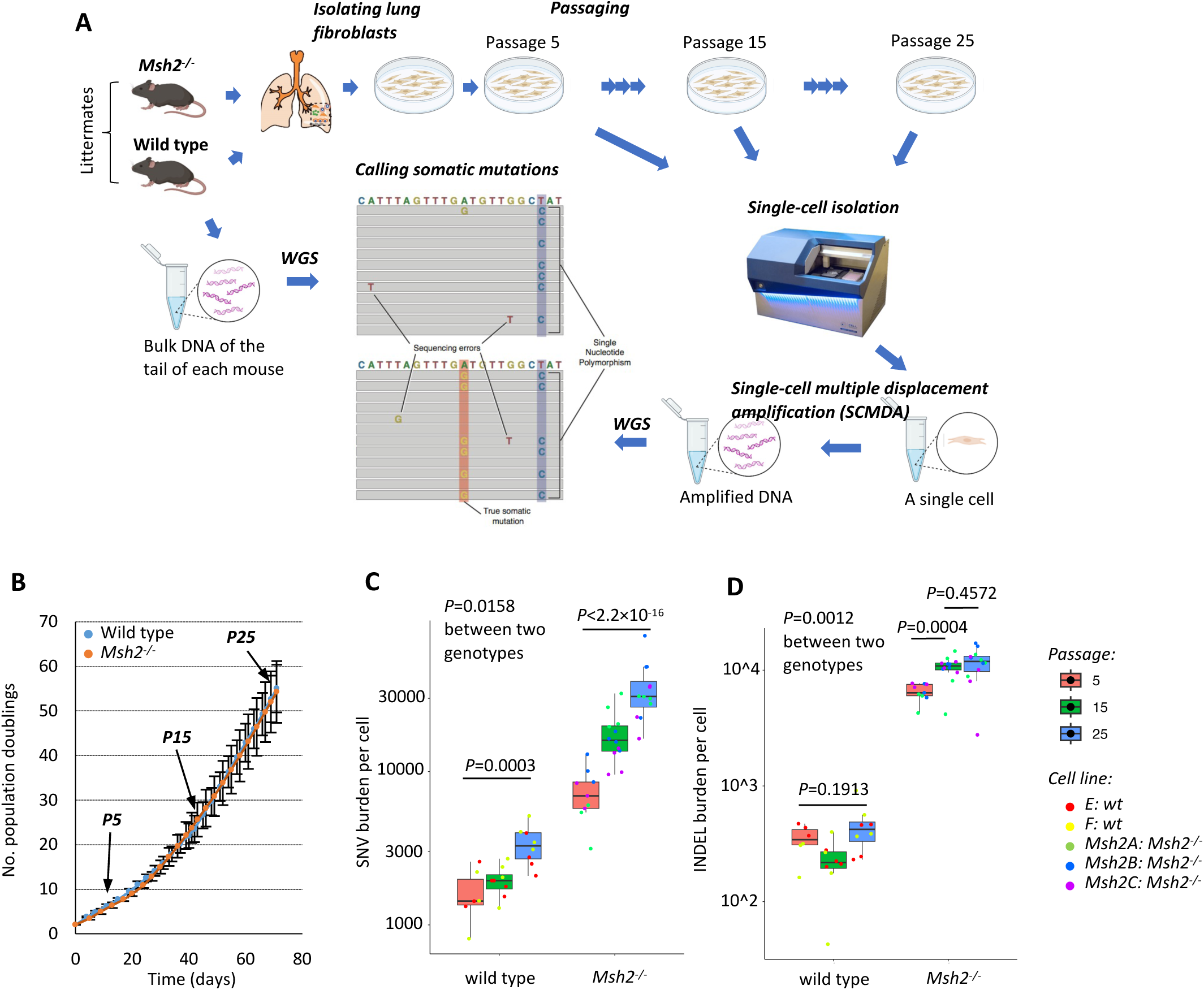
Study design, cell growth, and mutation burden. (**A**) A schematic illustration of the study design. We isolated lung fibroblasts of Msh2-/- and wild-type mice, and cultured them for 25 passages. De novo mutations in fibroblasts in passages 5, 15, and 25 of the cell strains obtained from different animal subjects were analyzed using single-cell whole-genome sequencing and compared to bulk whole-genome sequencing of the tails of the corresponding animals. (**B**) Cell growth during passaging. Error bars present s.d. (**C**) SNV burden and (**D**) INDEL burden per cell on log scales. Each data point presents a cell. *P* values were estimated using linear mixed effects models, two-sided using the “nlme” package of R. Boxplot elements are defined as follows: center line indicates median, box limits indicate upper and lower quartiles, and whiskers indicate 1.5× interquartile range.

## Results

### Somatic mutation burden in *Msh2^-/-^* mouse fibroblasts

Mice nullizygous for the *Msh2* gene, were generated and backcrossed into C57BL/6 as described previously (Smits et al., 2000). Their genotypes were validated using Polymerase Chain Reaction (PCR) of the DNA extracted from their tails (**Fig. S1**). Lung fibroblasts isolated from three *Msh2^-/-^*mice (4-5 months) and four wild-type mice, i.e., two wild-type littermates (4-5 months) and two additional, non-littermate wild-type mice (C57BL/6, 6 months), were cultured for 25 passages up to a total of 62 population doublings (**Methods**). As shown in **Fig. 1B**, growth rates of the three *Msh2^-/-^* and four wild-type fibroblast strains are almost identical, with no morphologic evidence for neoplastic transformation.

To quantitively analyze somatic mutation burden, we performed single-cell whole genome sequencing (scWGS) on 55 single cells at passages 5, 15, and 25 (denoted as P5, P15, and P25, respectively) of the three *Msh2*^-/-^ cell strains and the two wild-type littermate cell strains (**Fig. 1A**; **Methods**). Of note, the Single-Cell Multiple Displacement Amplification (SCMDA) and variant calling procedure (SCcaller) have been designed to avoid artificial mutations, previously the major problem in somatic mutation analysis (Dong *et al*., 2017; Zhang et al., 2023). For each cell strain, we also performed whole-genome sequencing of tail DNA from the same mice to identify germline polymorphisms, which were filtered out in calling *de novo* somatic mutations from the single cells. Depth of sequencing reached on average of 27.5x and 21.4x per sample for single cells and bulk DNAs, respectively (**Table S1**), to ensure that mutations could be identified accurately.

From the scWGS data on the 5 cell strains, we identified a total of 192,933 *de novo* mutations, including 147,955 SNVs and 44,978 INDELs, which was sufficient for analyzing mutation burden, spectrum, and distribution across the genome, especially for the *Msh2^-/-^* strains because of their high mutation frequencies (below). After correcting for sensitivity of variant calling and genome coverage (**Table S2**), we found that, as expected, *Msh2^-/-^* cells had a significantly higher SNV burden than wild type cells across all passages (*P*=0.0158, linear mixed effects model, two-sided). In wild type cells SNV burden increased with passage number in fibroblasts from 1,632±646 per cell (avg.±s.d.; P5) to 3,382±984 per cell (P25) in the wild type cells (*P*=0.0003, linear mixed effects model, two-sided), i.e., a 2-fold increase (**Figs. 1C** and **S2A**), which correspond to a mutation rate of ∼6.5x10^-9^ per bp per mitosis, almost the same as we estimated earlier for mouse primary fibroblasts (8.1x10^-9^ per bp per mitosis) (Milholland et al., 2017). In the *Msh2^-/-^* cells SNV burden increased from 7,475±2,902 per cell (P5) to 35,456±16,142 per cell (P25) (*P*<2.2×10^-16^, linear mixed effects model, two-sided), i.e., a 4.7-fold increase. There was no sign of a plateau between P5 and P25, not even in the *Msh2^-/-^* cells after acquiring tens of thousands of SNVs per cell. At P5, SNV burden in *Msh2^-/-^* cells was more than 4-fold higher than in the same cells from its littermate controls. Since we did not compare cells at different stages of embryonic development, we do not know how many more somatic mutations were present in the *Msh2^-/-^* mice from embryogenesis to early adulthood as compared to control mice, but it is safe to say that the original estimates based on reporter genes have been seriously overstated, i.e., 35- 550 mutations per 10^-5^ bp, corresponding to 1-15x10^6^ mutations per cell) (Hegan *et al*., 2006).

INDELs showed a different pattern of accumulation during passaging than SNVs (**Figs. 1D** and **S2B**). As expected, *Msh2^-/-^* cells had a significantly higher INDEL burden than the wild type cells across all passages (*P*=0.0012, linear mixed effects model, two-sided). However, INDEL burden during passaging only increased by 1.6-fold in the *Msh2^-/-^* cells between P5 and P15 (6,514±1,119 and 10,502±2,563 INDELs per *Msh2^-/-^* cell for P5 and P15, respectively; *P*=0.0004, linear mixed effects model, two-sided), but not between P15 and P25 (10,502±2,563 and 11,472±3,808 INDELs per *Msh2^-/-^* cells for P15 and P25 separately; *P*=0.4572, linear mixed effects model, two-sided). In cells from the littermate controls, no significant increase was observed during passaging (344±51 and 454±216 INDELs per cell for P5 and P25 separately; *P*=0.1913, linear mixed effects model, two-sided). These results indicate that INDEL tolerance reaches an upper limit in both wild type and *Msh2^-/-^* cells, but earlier in the control cells, possibly because most INDELs due to *Msh2^-/-^* are located in mononucleotide repeat sequences (see INDEL signature analysis below). Overall, these results indicate that the observed high numbers of SNVs or INDELs do not adversely affect growth rate of primary fibroblasts.

### Selection against damaging mutations

These results appear to suggest that increased burden of somatic mutations per se, i.e., without clonal amplification, do not cause cellular degeneration and death. Indeed, somatic mutation burden in tissues of aged humans or mice of the types of mutations analyzed here, never reach levels as observed in the MMR-deficient cells (Ren et al., 2022). However, while during *in vivo* aging selection against mutations that affect cellular function is difficult to envision, primary fibroblasts expanded *in vitro* offer an immediate mechanism of avoiding adverse somatic mutations by selection against mutations causing growth inhibition. Also, INDELs are generally more damaging than SNVs, many of which are synonymous and have no impact at all. To address the different impact of INDELs and SNVs in *Msh2^-/-^* and control cells during passaging we performed three comparisons as follows.

First, to test if the selection against INDELs is significantly stronger than the selection against SNVs, we calculated the ratio of INDEL burden to SNV burden for each single cell. As shown in **Fig. 2A**, INDEL-to-SNV ratio decreases significantly in fibroblasts of both genotypes: from 0.24±0.12 (P5) to 0.13±0.03 (P25) in the wild type cells (*P*=0.0223, linear mixed effects model, two-sided), i.e., a 1.8-fold decrease; and from 0.97±0.29 (P5) to 0.34±0.12 per cell (P25) in the *Msh2^-/-^* cells (*P*<2.2×10^-16^, linear mixed effects model, two-sided), i.e., a 2.8-fold decrease. These results indicate negative selection against INDELs during passaging in cells of both genotypes.

**Figure 2.**
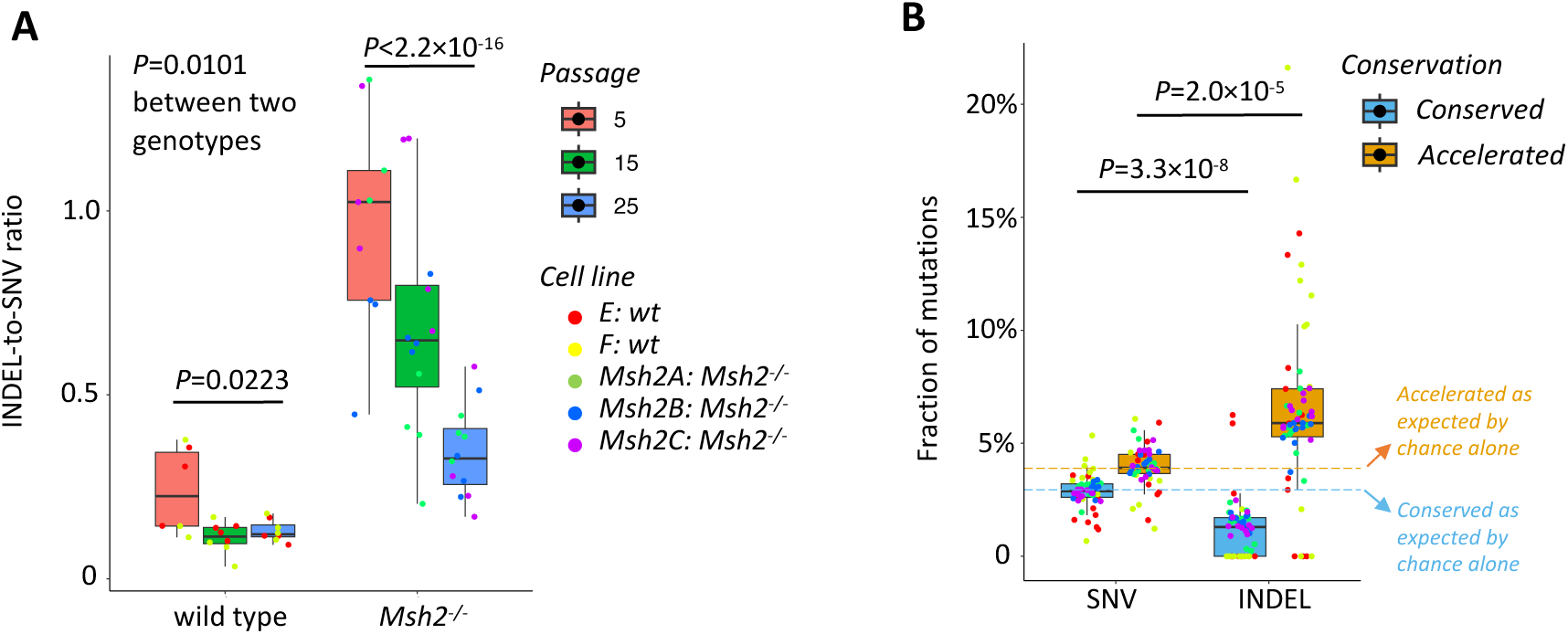
Selection pressures against INDELs. (**A**) The ratio of the number of INDELs to the number of SNVs per cell. (**B**) The fractions of mutations (SNVs and INDELs combined) at evolutionarily conserved and accelerated sites out of total mutations per cell. The fractions of SNVs and INDELs at conserved and accelerated sites by chance alone were estimated based on randomly generated mutations using SigProfilerSimulator (Bergstrom et al., 2020) – we randomly generated the same number of SNVs and INDELs as the observed numbers with also the same mutation signature, performed the same analysis of their conservation scores, and repeated the above two steps 2,000 times to reach stable estimations. Because there is no difference between the values of SNVs and INDELs expected by chance alone, we merged into two single values as indicated by the two dashed lines (for conserved and accelerated sites separately). Boxplot elements are defined as: center line indicates median, box limits indicate upper and lower quartiles, and whiskers indicate 1.5× interquartile range.

Second, to evaluate possible negative selection for both INDELS and SNVs, we utilized phyloP scores (Pollard et al., 2010; Siepel et al., 2005), with a positive score indicating conservation, i.e., slower evolution than expected, and a negative score indicating acceleration, i.e., faster evolution than expected. We obtained phyloP scores for all bases of the mouse reference genome from the UCSC genome browser (Lee et al., 2022). We then defined mutations at evolutionarily conserved sites as those with a phyloP score >0, its original *P* value <0.05, and percentile of the phyloP score of the mutated site as compared to the phyloP scores of its ±500 flanking bases >95% (which is to avoid a potential difference in genome coverage). Mutations at evolutionarily accelerated sites were defined by a phyloP score <0, its original *P* value <0.05, and percentile of the phyloP score of the mutated site as compared to the phyloP scores of its ±500 flanking bases<5%.

For both SNVs and INDELs in both wild-type and *Msh2^-/-^*cells, the fraction of mutations at an evolutionarily conserved site was substantially lower than that at an accelerated site (**Fig. 2B**). However, compared to mutations randomly sampled from the genome, we found that the fractions of SNVs at both conserved and accelerated sites were as expected by chance alone, while the fractions of INDELs were substantially different from the random sampling. A significantly smaller fraction of INDELs (1.2%±1.3%) was observed at a conserved site than SNVs (2.9%±0.8%; *P*=3.3×10^-8^, paired Wilcoxon signed-rank tests, two-sided) or expected based on chance alone. By contrast, a greater fraction of INDELs was found at an accelerated site than SNVs (6.6%±4.0% and 4.0%±0.9% for INDELs and SNVs respectively; *P*=2.0×10^-5^, paired Wilcoxon signed-rank tests, two-sided) or as expected by chance alone. Of note, in 77% of wild type cells we did not observe any INDELs at a conserved site. During passaging, no significant change was observed between SNVs and INDELs at accelerated and conserved sites in cells of the two genotypes (linear mixed effects models, two-sided; **Figs. S3A-D**) with two exceptions: a marginal increase of INDELs at conserved sites in *Msh2^-/-^* cells (*P*=0.0455, i.e., no longer significant if adjusting for multiple testing; **Fig. S3C**); and a significant decrease of SNVs at accelerated sites in *Msh2^-/-^*cells (*P*=0.0011; **Fig. S3B**). Overall, these results indicate negative selection at evolutionarily conserved sites for INDELs during passaging, but not for SNVs.

Finally, we performed bulk RNA sequencing of each fibroblast cell strain to determine genes that are transcriptionally active (**Methods**). Using mutation annotation by ANNOVAR (Wang et al., 2010; Yang and Wang, 2015), we then analyzed mutations that alter protein coding sequences of transcriptionally active genes (**Table S3**). We calculated the ratio of nonsynonymous to synonymous SNVs in the two genotypes during passaging and found that this ratio remains approximately the same and shows no significant difference from the ratios expected by chance alone (**Figs. 3A-B**), suggesting a lack of negative selection. However, significantly less frameshifting INDELs than expected by chance alone were found in these cells during passaging (0.05±0.21 per cell and 3.7±2.6 per cell for wild type and *Msh2^-/-^* cells separately), as well as significantly less stop-gain SNVs (0.14±0.47 per cell and 1.0±1.5 per cell for separately), or stop-loss SNVs (0±0 per cell and 0.03±0.17 per cell for wild type and *Msh2^-/-^* cells separately) (**Figs. 3 C, D, F, G, I & J**). This is in keeping with our previous observations that in human B cells from aged human subjects on average less than one loss-of-function mutation (including stop-gain, stop-loss, and splicing alteration) per cell was observed (Zhang *et al*., 2019). The absence of MMR in the *Msh2*^-/-^ cells rules out preferential protection of actively transcribed genes (Huang and Li, 2018) as a mechanism to explain the observed lower rates of deleterious mutations. Yet, pre-replication, transcription-coupled repair (TCR) of DNA damage could still explain these results, at least in part (Georgakopoulos-Soares et al., 2020). To test whether the reduced observed-to-expected ratios of loss-of-function mutations are due to negative selection or increased DNA repair activity, we estimated the ratio of each type of loss-of-function mutation to synonymous mutations and compared the ratios to those expected by chance alone. As shown in **Figs. 3 E, H & K**, most of the ratios are significantly smaller than expected by chance alone, indicating that the limited numbers of loss-of-function mutations are a result of negative selection, and not due to increased DNA repair in transcribed regions.

**Figure 3.**
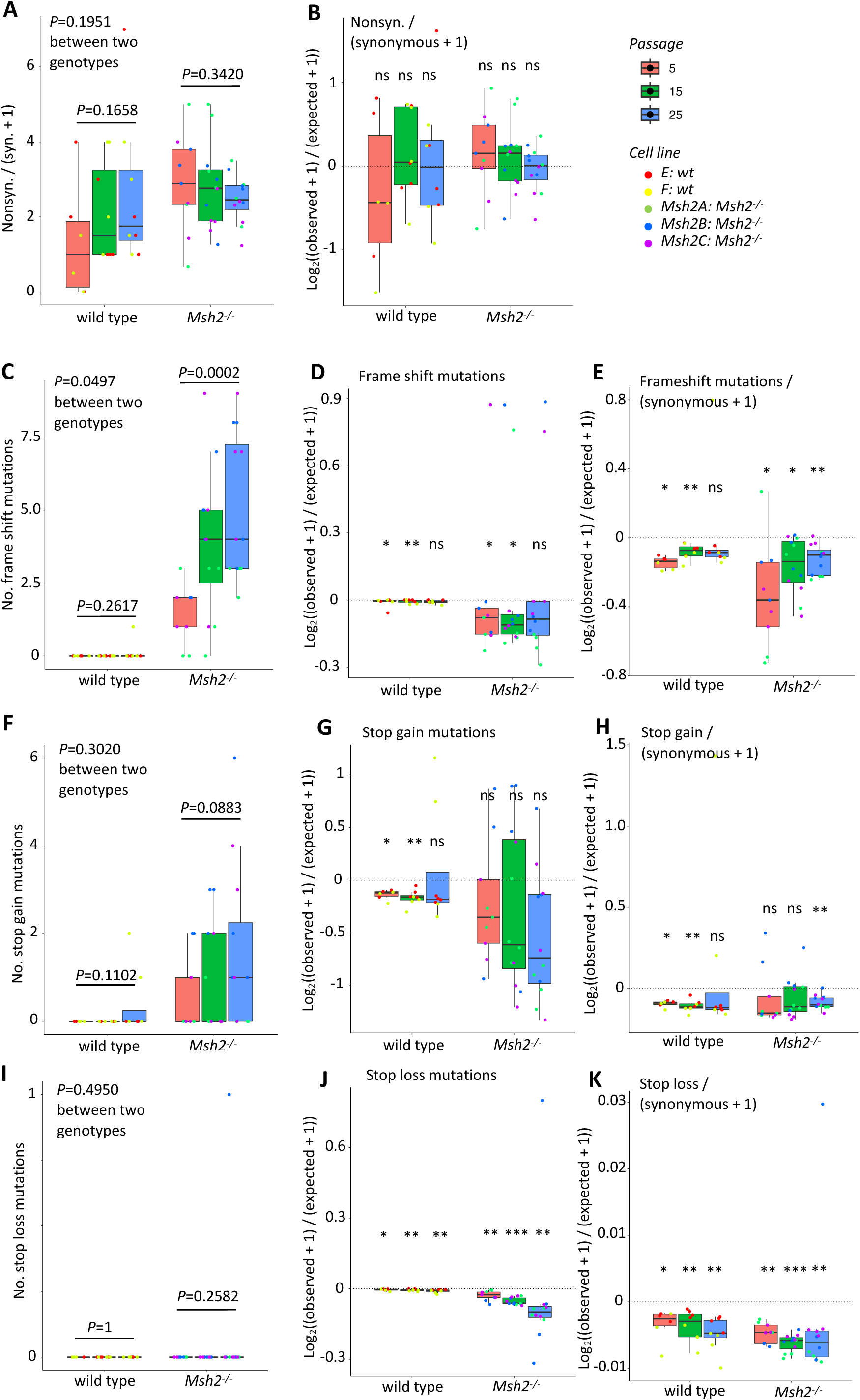
Selection pressures against damaging mutations. (**A**) The ratio of the number of nonsynonymous mutations to synonymous mutations. We added 1 to the denominator values to avoid potential 0. (**C, F & I**) The numbers of frameshifting, stop-gain, and stop-loss mutations per cell. (**D, G & J**) The numbers of observed frameshifting, stop-gain, and stop-loss mutations vs. their corresponding numbers expected by chance alone (in log2-transformed ratios). (**B, E, H & K**) The observed ratios of the numbers of nonsynonymous, frameshifting, stop-gain, and stop-loss mutations to the numbers of synonymous mutations vs their corresponding ratios expected by chance alone. To estimate the number of mutations expected by chance alone, we first used SigProfilerSimulator (Bergstrom *et al*., 2020) to randomly generate the same number of SNVs and INDELs as the observed numbers with also the same mutation signature, then annotated the artificial mutations with ANNOVAR (Yang and Wang, 2015) to determine the number of mutations in each functional category, and finally repeated the above two steps 2,000 times to reach stable estimations. Each dot presents a cell. *P* values in **A, B, C, D, F, & H** were estimated using linear mixed effects models, two-sided. In E, G, & I, “ns”, “*”, “**”, and “***” represents *P* values >0.05, <0.05, <0.01, and <0.001, separately, which were estimated using binomial tests, two sided. Boxplot elements are defined as: center line indicates median, box limits indicate upper and lower quartiles, and whiskers indicate 1.5× interquartile range.

### Each *Msh2^-/-^* cell strain acquires common and unique mutational signatures during passaging

As shown in studies of human cancers, mutational spectra and signatures suggest specific factors that drive mutagenesis, e.g., oxidative damage, radiation (Alexandrov et al., 2020; Alexandrov et al., 2013). However, connection between mutation signatures and causal factors are often derived computationally. In this study, we had an opportunity to test if passaging and DNA mismatch repair deficiency indeed causes the mutational signatures inferred from human cancers.

First, we compared SNV spectra between the cell strains. As expected, *Msh2^-/-^* cells are substantially different from wild-type cells with more C>T and T>C mutations (**Fig. S4A**). However, we noticed substantial variation between the three *Msh2^-/-^* cell strains: the Msh2A cell strain acquired more T>C mutations, the Msh2C cell strain acquired more C>T mutations, and the Msh2B cell strain was in between (**Fig. 4A**). Of note, their unique mutational spectra became more obvious during passaging (**Fig. S4B**).

**Figure 4.**
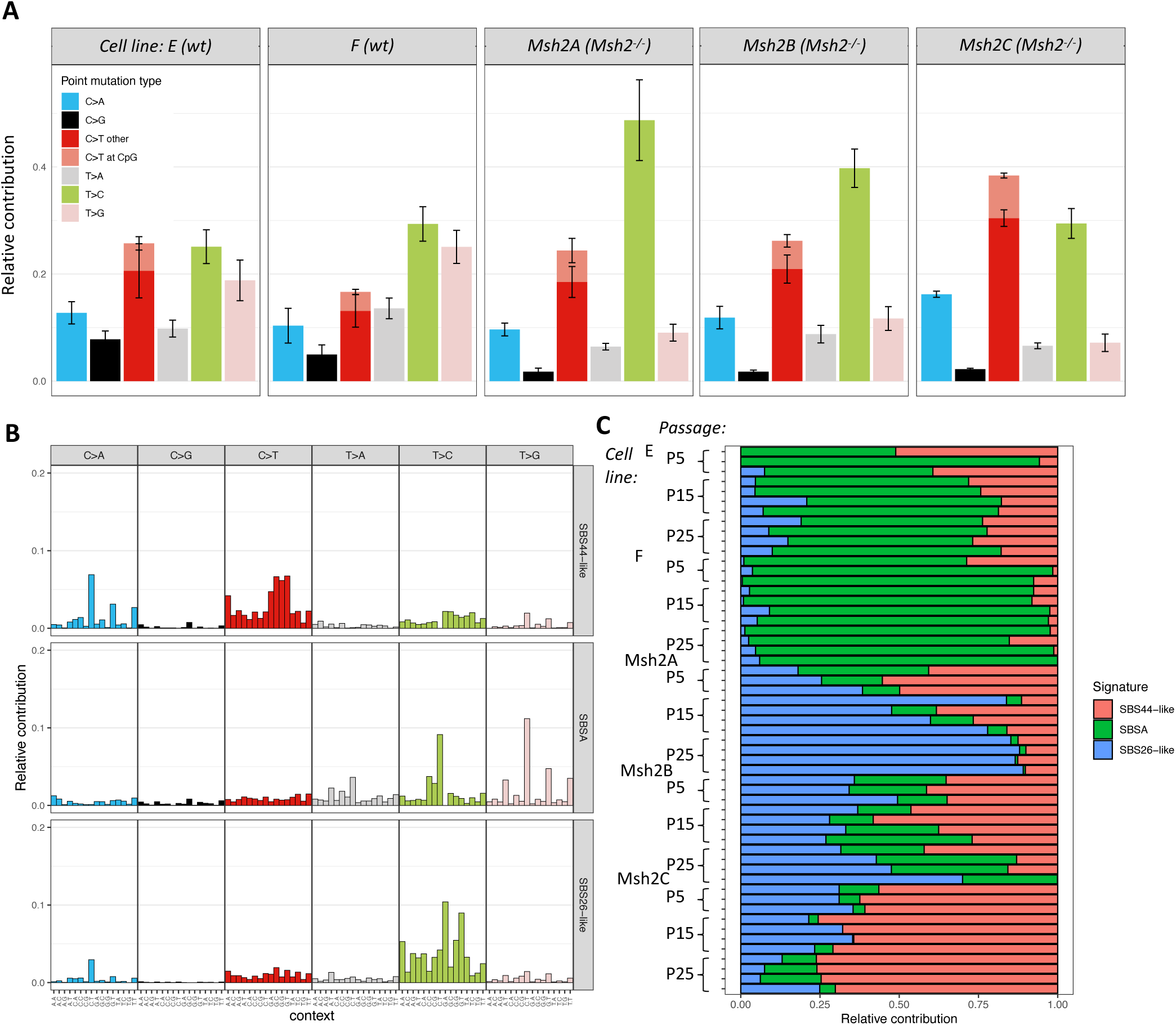
SNV spectra and signatures. (**A**) SNV spectra of each cell strain. Error bars present s.d. (**B**) Three SNV signatures of the fibroblasts identified by *de novo* signature extraction using the “MutationalPatterns” package of R(Blokzijl *et al*., 2018). (**C**) Contribution of each SNV signature to the total SNVs per cell.

Then, we performed SNV signature analyses in two ways, both using the “MutationalPatterns” package of R (Blokzijl et al., 2018). First, we performed *de novo* signature extraction, and identified three signatures (**Fig. 4B**). Using a cosine correlation cutoff at 0.85 with known mutational signatures of human cancers reported in the COSMIC database (Alexandrov *et al*., 2020), we labeled the three signatures as SBS-A (no similar cancer signature was found), SBS26- like (positively correlated with the COSMIC Single Base Substitution signature #26), and SBS44-like signatures. The SBS26-like signature dominates mutations in the Msh2A cell strain and its fraction out of all mutations increases with passaging, while the SBS44-like signature is more dominant in the Msh2C cell strain (**Fig. 4C**). Of note, both SBS26 and SBS44 signatures in tumors have been suggested to be the result of DNA mismatch repair deficiency (Alexandrov *et al*., 2020). The SBS-A signature, which was not reported in the COSMIC database, contributes to most mutations in the wild type cells (**Fig. 4C**) and is likely a result of replication errors. However, SBS-A (characterized by N<u>T</u>T>N<u>G</u>T or N<u>C</u>T mutations; **Fig. 4B**) is very different from the SBS1 signature (characterized by N<u>C</u>G>N<u>T</u>G mutations (Alexandrov *et al*., 2020)) in human tumors, which has been associated with cell division.

Second, we refitted COSMIC signatures to the mutations that we observed. When doing that we found another DNA mismatch repair signature, i.e., SBS21, in the *Msh2^-/-^* cell strains, but the differences between the *Msh2^-/-^* cell strains remained (**Fig. S5**). Together, despite confirming that MMR deficiency can indeed cause the corresponding signatures found in human cancers, these results indicate that a single factor, i.e., *Msh2*-deficiency, can result in different mutational signatures.

For INDELs, we also performed signature extraction, and identified two signatures: an ID2-like signature (positively correlated with the COSMIC small Insertion and Deletion signature #2), which is characterized as a single-base T deletion in repetitive T sequences, and another new signature, termed IDA, which does not correlate with a COSMIC signature (**Fig. S6A**). IDA was mostly found in our wild-type control cells (**Fig. S6B**) and is characterized by either insertion or deletion at repeat regions of multiple homopolymers or repeat units. The ID2-like signature, mostly single base deletions in a long homopolymer of thymines, was predominantly found in our *Msh2^-/-^* cell strains (**Fig. S6B**). The ID2 signature in human cancers is suggested to be caused by slippage during DNA replication of the template DNA strand and is often found in DNA mismatch repair deficient tumors (Alexandrov *et al*., 2020). Of note, in the COSMIC database, another INDEL signature, ID7, characterized by 1-bp deletions at homopolymers of both cytosines and thymine and suggested to be a result of MMR deficiency in humans, was not observed here.

### Hotspots and overlap of mutations

We then tested for mutational hotspots (for SNVs and INDELs together) in the mouse genome by using the “ClusteredMutations” package in R (Lora, 2016). A substantial number of mutational hotspots were observed in both WT and *Msh2^-/-^* fibroblasts, but significantly more in the latter (**Fig. 5A**). Surprisingly, mutational hotspots were so obvious, even in wild-type cells, that we could identify them for each individual cell, while in our previous study of human lymphocytes we had to pool mutations observed in tens of cells to discover significant mutational hotspots (Zhang *et al*., 2019). We then used a rainfall plot to visualize the distribution of the mutational hotspots across the genome. Again, different cell strains showed substantially different patterns (**Fig. 5B**). The Msh2A strain continuously gained additional mutational hotspots at the end of chromosome 17, while in the Msh2B cell strain, which showed the highest number of mutational hotspots, these spread across the entire reference genome during passaging. Two “super- hotspots” are worth noticing. One is at chr17:86,631,535-90,041,858 bp, found exclusively in the Msh2A cell strain. Interestingly, *Msh2* and *Msh6* genes locate in this region along with over 20 other genes, but all mutations in the hotspots at this region locate at intergenic sequences. The other super-hotspot was found at chr1:170,941,871-170,943,280 bp and was observed in four of the five cell strains (two WT and two *Msh2*^-/-^), but not in the Msh2A strain. This region is entirely intergenic and is part of a LTR repeat element.

**Figure 5.**
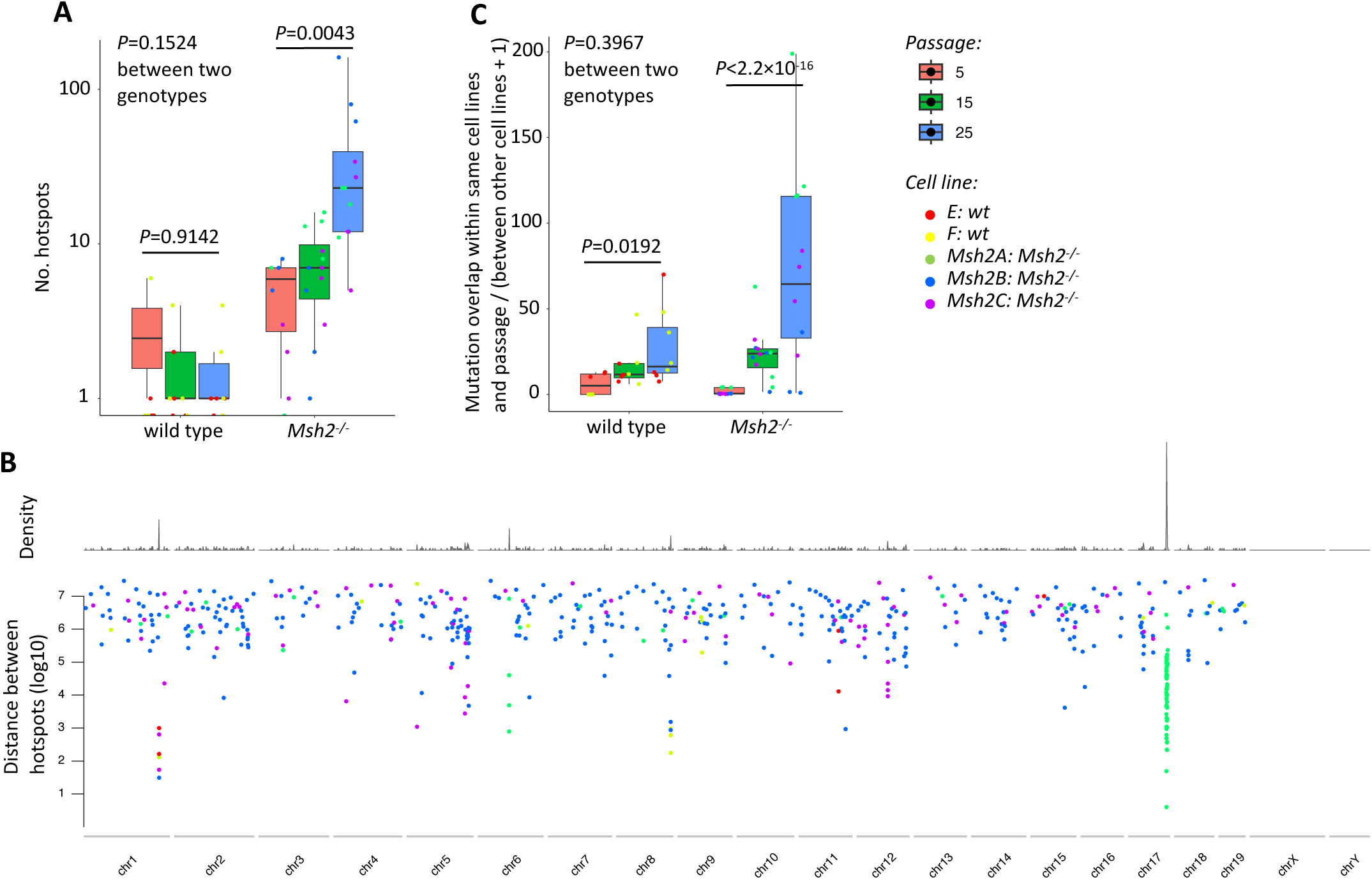
Mutational hotspots and overlap. (**A**) The number of mutational hotspots (SNV and INDELs combined) per cell. (**B**) The ratio of the number of overlapping mutations among cells of the same passage and same cell strain (i.e., animal) to the number of overlapping mutations among all cells of all strain. We added 1 to the denominator values to avoid potential 0. Each data point presents a cell. *P* values were estimated using linear mixed effects models, two-sided. Boxplot elements are defined as follows: center line indicates median, box limits indicate upper and lower quartiles, and whiskers indicate 1.5× interquartile range. (**C**) A rainfall plot of the distribution of mutational hotspots across the genome. The plot was generated using the “karyoploteR” package of R(Gel and Serra, 2017). Each data point presents a mutational hotspot observed within a single cell.

Why would each *Msh2*^-/-^ cell strain develop its own unique pattern of mutational hotspots? It is possible that substantial clonal expansion occurred during passaging, and each cell strain was eventually dominated by different clones. To test this, we calculated for each cell in each cell strain (of both WT and *Msh2*^-/-^) the ratio of (a) the mutations overlapping with mutations in other cells of the same passage and cell strain to (b) the mutations found to overlap in all cells of all cell strains. A higher ratio indicates more clonal expansion. As shown in **Figs. 5C** and **S7**, ratios increase dramatically during passaging in cell strains of both genotypes: from 6.0±6.6 (P5) to 27.3±22.1 (P25) in wild type cells (*P*=0.0192, linear mixed effects model, two-sided); and from 1.7±1.9 (P5) to 71.9±58.7 (P25) in *Msh2^-/-^* cells (*P*<2.2×10^-16^, linear mixed effects model, two-sided). Although the difference between cells of the two genotypes was not statistically significant (*P*=0.3967, linear mixed effects model, two-sided), likely due to large cell-to-cell variations, the increase in *Msh2^-/-^* cells is substantially higher (a 42-fold increase from P5 to P25) than in the wild type cells (a 4.6-fold increase). These results confirm the occurrence of substantial clonal expansion during passaging in cells of both genotypes, with different cell strains taken over by different clones. This process is a likely cause of the different mutational signatures and hotspots observed in different cell strains. These results also suggest strong positive selection of specific cell lineages in the different cell strains, which is frequently observed in tumor cells (Martincorena et al., 2017).

## Discussion

With the emergence of advanced high-throughput sequencing methods, including high-accuracy single-cell and single-molecule methods, increased insights are now being obtained in somatic rather than germline mutations as a possible cause of human genetic disease and aging (Mustjoki and Young, 2021; Vijg and Dong, 2020). Mutation frequency in somatic cells and tissues appeared to be 1-2 orders of magnitude higher than germline mutation frequency (Milholland *et al*., 2017). This is in keeping with the disposable soma theory of aging, which states that reproduction is prioritized over somatic maintenance (Kirkwood, 1977). This idea is in line with the observed correlation of somatic maintenance and species-specific life span (Hart and Setlow, 1974). Indeed, we and others recently showed that somatic mutation rate is inversely correlated with species-specific life span (Cagan et al., 2022; Zhang et al., 2021).

Recent findings that somatic mutation burden increases with age in different human tissues (Ren *et al*., 2022) supports a possible causal role of somatic mutations in the aging process. Indeed, clonally amplified somatic mutations, which are relatively easy to detect by high-depth sequencing, have now been shown to be a cause of a large number of human diseases other than cancer (Erickson, 2010; Mustjoki and Young, 2021). However, what remains unclear is if increased somatic mutation burden per se can cause cellular degeneration and death. In this respect, a key question is if random somatic mutations can rise to a level high enough to infringe on the integrity of the gene regulatory pathways that provide function to the specialized somatic cells in the human body. Here we present mutation accumulation data for a simplified cell culture model in the form of mouse primary fibroblasts with mutations continuously generated through a defect in DNA mismatch repair.

The first conclusions that can be drawn based on our data is that somatic SNVs can accumulate to levels at least 6 times as high as observed in human postmitotic tissues from aged subjects (Brazhnik *et al*., 2020; Lodato *et al*., 2018). Our finding that these high numbers of random mutations have no significant effects on growth rate seems to rule out a causal role of somatic mutations in aging. However, in contrast to the situation during normal aging, cell culture systems are subject to selection against deleterious mutations affecting growth. We found ample evidence for such selection in all fibroblast strains studied, including the control, wildtype strains. First, among SNVs we found significant negative selection against stop-loss and stop- gain mutations. Second, while SNV burden never reached plateau levels up until a population doubling level (PDL) of 50-60 (i.e., P25, **Fig. 1C**), INDEL burden did not increase in controls and no longer increased after 20-30 PDL (i.e., P15) in the Msh2-deficient cells. These observations are different from mutations in human tumors, in which positive selection has been shown to outweigh negative selection (Martincorena *et al*., 2017).

Of note, in mitotically active human B lymphocytes we previously found the rate of age-related SNV accumulation in the about 10% functionally active part of the genome to be only half of the genome-wide average (Zhang *et al*., 2019). Yet, except for loss-of-function SNVs, which do not increase with age in human lymphocytes, the number of potentially functional SNVs still accumulated with age, even in subjects in their 80s- or 90s (Zhang *et al*., 2019).

In addition to the evidence for direct selection against deleterious mutations, most notably INDELS, we also found evidence for widespread mutational hotspots and significant clonal expansion. Both differed between the cell strains studied, gradually leading to unique populations in each strain. Together with direct selection against deleterious mutations, such mutational evolution could be responsible for maintaining normal growth rate even after acquiring tens of thousands of SNVs and almost 10,000 INDELS in the Msh2-deficient cells.

The fact that somatic mutations, either spontaneous or driven by the MMR defect, show such dramatic evolutionary dynamism in culture, strongly suggests they have functional consequences. If they would be completely neutral, none of these effects would be expected to occur. However, with some possible exceptions (e.g., the lymphoid and intestinal systems) adult tissues have limited options for negative selection since most are not mitotically active. While the observation of clonally amplified mutations in virtually all tissues, most notably clonal hematopoiesis (Jaiswal and Ebert, 2019), demonstrate positive selection for a growth or survival advantage, we now show that negative selection may occur as well. In the absence of such selection it is conceivable that random mutations at the levels observed in aged subjects will gradually impair cellular function in somatic cells (Vijg and Dong, 2020).

At least one limitation of our current study should be mentioned, which is the driver of the high level of somatic mutagenesis itself. MMR deficiency does not elevate all categories of mutations equally and it can be argued that the most impactful mutations, including genome structural variation, are not significantly elevated at all. Indeed, this could be one of the reasons of a lack of premature aging in MMR-deficient mice or humans (Robinson et al., 2021). Another reason could simply be the lack of detailed analysis of premature aging in MMR-deficient mice or humans, which usually die from cancer well before old age), which is not trivial (Franco et al., 2022).

In summary, our present data uncover the comprehensive landscape of somatic mutations in MMR-deficient mouse primary fibroblasts as compared to wildtype control cells passaged *in vitro*. The results show that the MMR-deficient cell populations maintain high growth rates in spite of an SNV burden of at least 30,000 mutations per cell, while INDEL burden reaches a plateau of about 10,000 per cell. Further analysis showed extensive somatic evolution, including negative selection to maintain growth rate, possibly by eliminating deleterious mutations. We conclude that in the absence of such selection options, deleterious effects of accumulating somatic mutations to the levels that have been observed *in vivo* is inevitable. Further research on cell populations that can be directly interrogated for a functional relationship between somatic mutation burden and specific cellular functions known to decline with age will provide a more definitive test of a causal relationship between somatic mutations and aging.

### Data Availability

Raw sequencing data will be submitted to the NCBI SRA database before the paper is accepted.

## Acknowledgements

L.Z. is supported by the American Federation for Aging Research (the Sagol Network GerOmic Award for Junior Faculty). J.V. is supported by NIH grants (P01 AG017242, P01 AG047200, P30 AG038072, U01 ES029519, U01 HL145560, and U19 AG056278). X.D. is supported by NIH grants (R00 AG056656, U19 AG056278, P01 HL160476, and P01 AI172501) and the Fesler-Lampert Chair for Aging Studies at the University of Minnesota.

## Materials and Methods

### Transgenic mice

Mice nullizygous for the *Msh2* gene, were generated and backcrossed into C57BL/6 as described previously(Smits *et al*., 2000). In this study, three *Msh2^-/-^* mice (4-5 months) and two of their wild- type littermates (4-5 months) were used. All procedures involving animals were approved by the Institutional Animal Care and Use Committee (IACUC) of Albert Einstein College of Medicine and performed in accordance with relevant guidelines and regulations.

### Bulk DNA extraction and genotyping

We extracted genomic DNA from tail of each mouse using the DNeasy Blood & Tissue Kit (Qiagen) following the manufacturer’s specifications. The concentrations of DNA were quantified using the Qubit High Sensitivity dsDNA Kit (Invitrogen Life Science) and the qualities of DNAs were evaluated with 1% agarose gel electrophoresis.

We validated the genotypes of the mouse strains by PCR genotyping using the genomic DNA as template. Each reaction contains 1 µl of gDNA (10ng/µl), 1.5 µl of 10x PCR buffer II (Roche), 1.5 µl of MgCl2 (25mM, Roche), 0.1 µl of Taq Gold (5U/ µl) and Primer A, B and C (The sequences of Primers are listed in **Fig. S1**). The total reaction volume of PCR is 12.5 µl. PCR conditions were 94 °C for 5 min; and 40 cycles 94 °C for 45 s, 55 °C for 1 min and 72 °C for 1 min; and 72 °C for 5 min. The PCR results were shown in the picture of 1% agarose gel electrophoresis (**Fig. S1**).

### Lung fibroblast isolation and passaging

Primary lung fibroblasts were isolated following a cell isolation protocol adapted from Seluanov *et al* (Seluanov et al., 2010). In brief, mouse lung was minced and incubated in DMEM F-12 medium with 0.13 unit/ml Liberase Blendzyme 3 and 1x penicillin/streptomycin at 37℃ for 40 min. Dissociated cells were washed, plated in cell culture dishes with complete DMEM F-12 medium, 15% FBS and cultured at 37℃, 5% CO_2_, 3% O_2_. When reaching confluence, cells were split and replated in EMEM medium supplemented with 15% FBS and 100 units/ml penicillin and streptomycin. Lung fibroblasts were purified by further passaging in the same medium.

From each subject, we passaged one cell strain. Cells from each cell strain were cultured and passaged in two 10cm-plate with EMEM supplemented with 15% FBS and 100 units/ml penicillin and streptomycin. The initial cell number was 0.5 or 1 million for each plate each passage. We counted cell numbers during passaging applying the Cellometer Auto T4 cell counter (Nexcelom), calculated cell population doublings based on the cell number of each cell strain and plotted the cell proliferation curve.

#### Single-cell isolation, whole-genome amplification, library preparation and sequencing

Single lung fibroblasts were isolated using the CellRaft AIR system (Cell Microsystems) according to the manufacturer’s instructions. Isolated single fibroblasts in 2.5 µl PBS were frozen immediately on dry ice and kept at -80℃ until amplification.

The isolated single fibroblasts were amplified using SCMDA as described(Dong *et al*., 2017). The amplicons were subjected to quality control using a locus dropout test(Milholland *et al*., 2017). Of those passing the quality control, three amplicons per mouse were subjected to library preparation and sequencing with 150-bp paired-end reads on an Illumina HiSeq X Ten sequencer (Novogene, Inc). Bulk DNAs extracted from tails of the same mice were sequenced without amplification and used for filtering out germline polymorphisms during variant calling as described(Dong *et al*., 2017).

#### Sequence alignment and mutation calling

Raw sequence reads were subject to quality control using FastQC (https://www.bioinformatics.babraham.ac.uk/projects/fastqc/), adaptor- and quality-trimmed using Trim Galore (https://www.bioinformatics.babraham.ac.uk/projects/trim_galore/), and aligned to reference genome mouse mm10 using bwa mem (Li and Durbin, 2009). PCR duplicates were removed using samtools (Li et al., 2009). The aligned reads were then INDEL-realigned and base-pair score quality recalibrated using GATK (McKenna et al., 2010). SNVs and INDELs observed in a cell but not presented in the corresponding bulk DNA of the tail were called by comparing the aligned sequences of the cell to the bulk using SCcaller (version 2.0) (Zhang *et al*., 2023): (i) from genomic regions covered with a minimum depth of 20x in both the cell and the bulk; (ii) with default parameters for SNVs; and (iii) requiring a variant calling quality ≥ 30 for INDELs. Mutation burden per cell were estimated based on the number of observed mutations adjusting coverage of the genome and variant calling sensitivity. For variant calling sensitivity in humans, we previously used the fraction of germline heterozygous mutations observed in the single cell, but the number of germline mutations is very limited in inbred mice. So instead, we used consistent values of sensitivity estimated from the scWGS data of fibroblasts of multiple 4- way-across mice and other rodent species as reported previously (**Table S2**) (Zhang *et al*., 2021).

#### Bulk RNA sequencing and data analysis

For each cell strain of different passages, total RNA was extracted using RNeasy Micro Kit (Qiagen) according to the manufacturer’s specification. The concentrations of RNA were quantified with Qubit RNA HS Assay Kit (Invitrogen Life Science) and the qualities of RNA were evaluated using bioanalyzer with Agilent RNA 6000 Pico Kit (Agilent Technologies). The qualified RNA samples (RIN≥7.0, OD260/280>2.0, concentration≥20ng/μl and volume≥20μl) were submitted to Novogene for library preparation and sequencing. The insert size of double-strand cDNA library is 250-300bp. The libraries of the RNA samples were sequenced on the Illumina Novaseq 6000, with 2×150 bp paired-end reads. The average sequencing amount of raw data of each library is 9.24 G bp.

Raw sequence reads were subject to quality control using FastQC (https://www.bioinformatics.babraham.ac.uk/projects/fastqc/), adaptor- and quality-trimmed using Trim Galore (https://www.bioinformatics.babraham.ac.uk/projects/trim_galore/), and aligned to reference transcriptome of mouse mm10 using STAR(Dobin et al., 2013). Gene expression levels were quantified using RSEM(Li and Dewey, 2011). Expressed protein coding genes were determined as those with an average transcript per million (TPM) value ≥ 1 across all samples.

---

**Figure S1.**
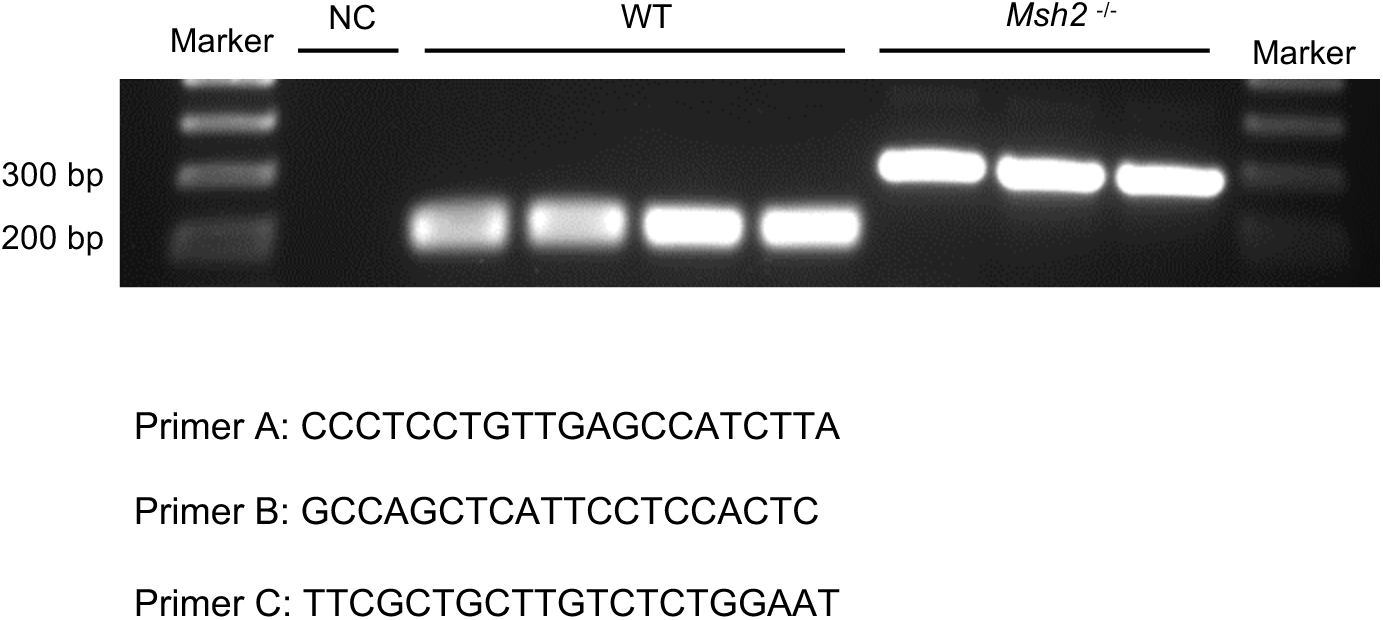
PCR genotyping. The genotypes of each mouse were validated using two pairs of primers, comprising three primers in total, with their sequences presented in this figure. Primer A was designed to align with the intron preceding exon 7 of the Msh2 gene, while primer C was situated in exon 7. Primer B was positioned in the PGK poly A cassette of the neomycin cassette, replacing an internal fragment that contains most of Msh2 exon 7. In wild-type mice, the A/C primer pair was used, resulting in an amplified PCR product of 189 bp. The wild-type group consisted of two C57BL/6 mice (shown in the two left lanes) and two wild-type littermates of the *Msh2^-/-^* mouse, E(wt) and F(wt) (shown in the two right lanes). For *Msh2^-/-^* mice, the A/B primer pair was used, producing a PCR product of 300 bp. The gel image of the *Msh2^-/-^* group displayed PCR results from three mice, Msh2A, Msh2B, and Msh2C (from left to right).

**Figure S2.**
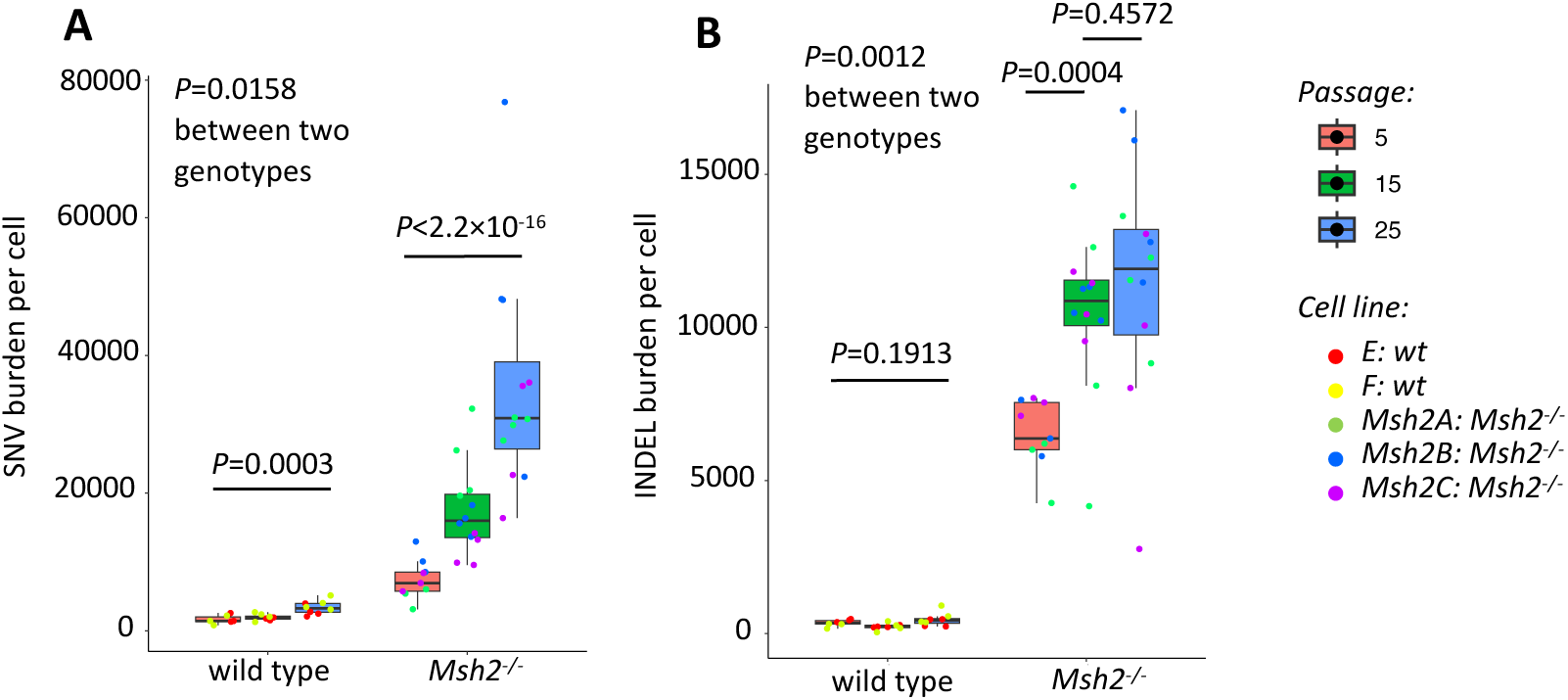
Mutation burden in linear scale. (**A**) SNV burden and (**B**) INDEL burden per cell in linear scale. Each data point presents a cell. *P* values were estimated using linear mixed effects models, two-sided. Boxplot elements are defined as follows: center line indicates median, box limits indicate upper and lower quartiles, and whiskers indicate 1.5× interquartile range.

**Figure S3.**
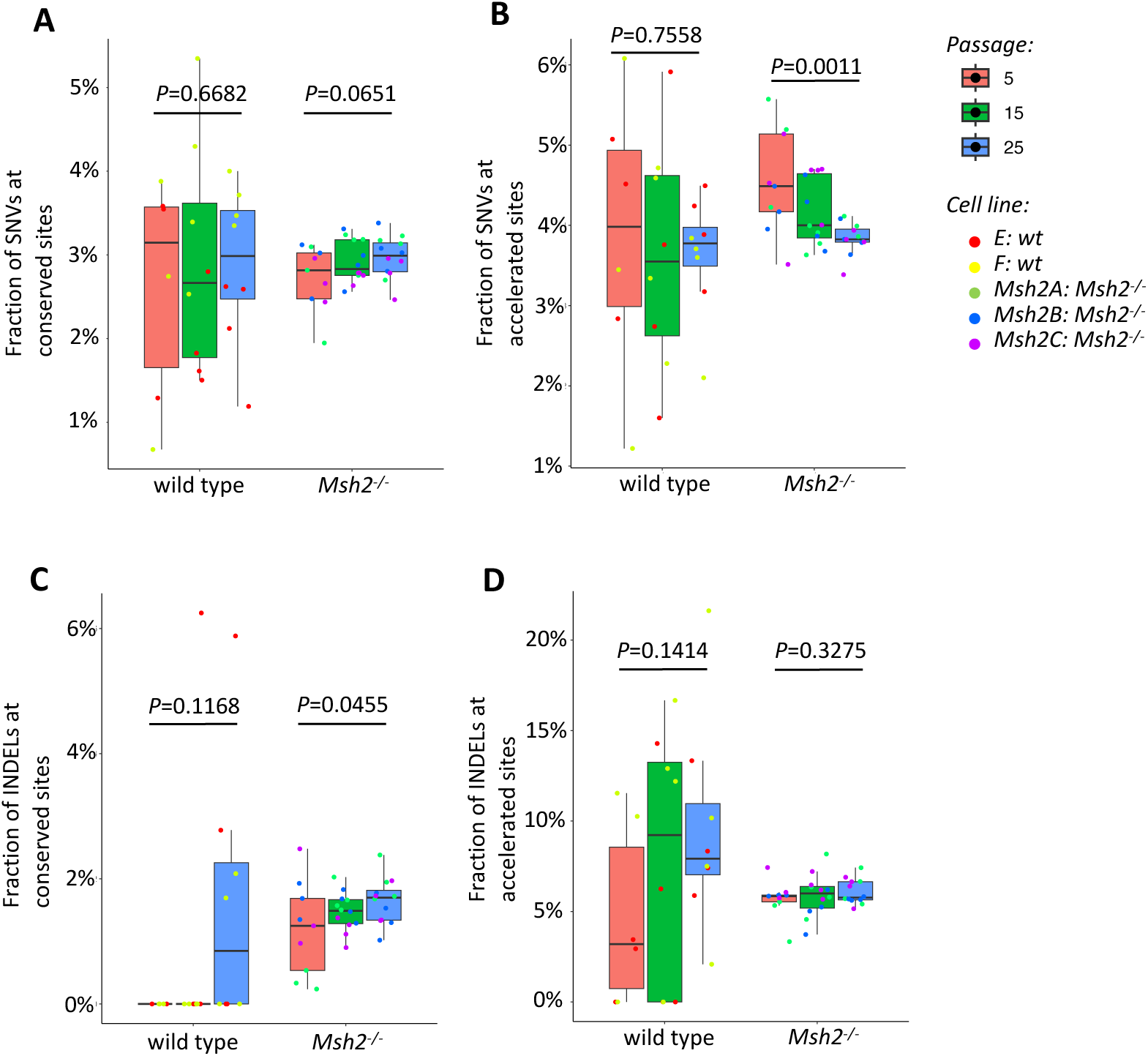
Mutation burden at evolutionarily conserved and active sites during passaging. The fractions of SNVs at evolutionarily (**A**) conserved and (**B**) accelerated sites out of total mutations per cell. The fractions of INDELs at evolutionarily (**C**) conserved and (**D**) accelerated sites out of total mutations per cell. Each data point presents a cell. *P* values were estimated using linear mixed effects models, two-sided. Boxplot elements are defined as: center line indicates median, box limits indicate upper and lower quartiles, and whiskers indicate 1.5× interquartile range.

**Figure S4.**
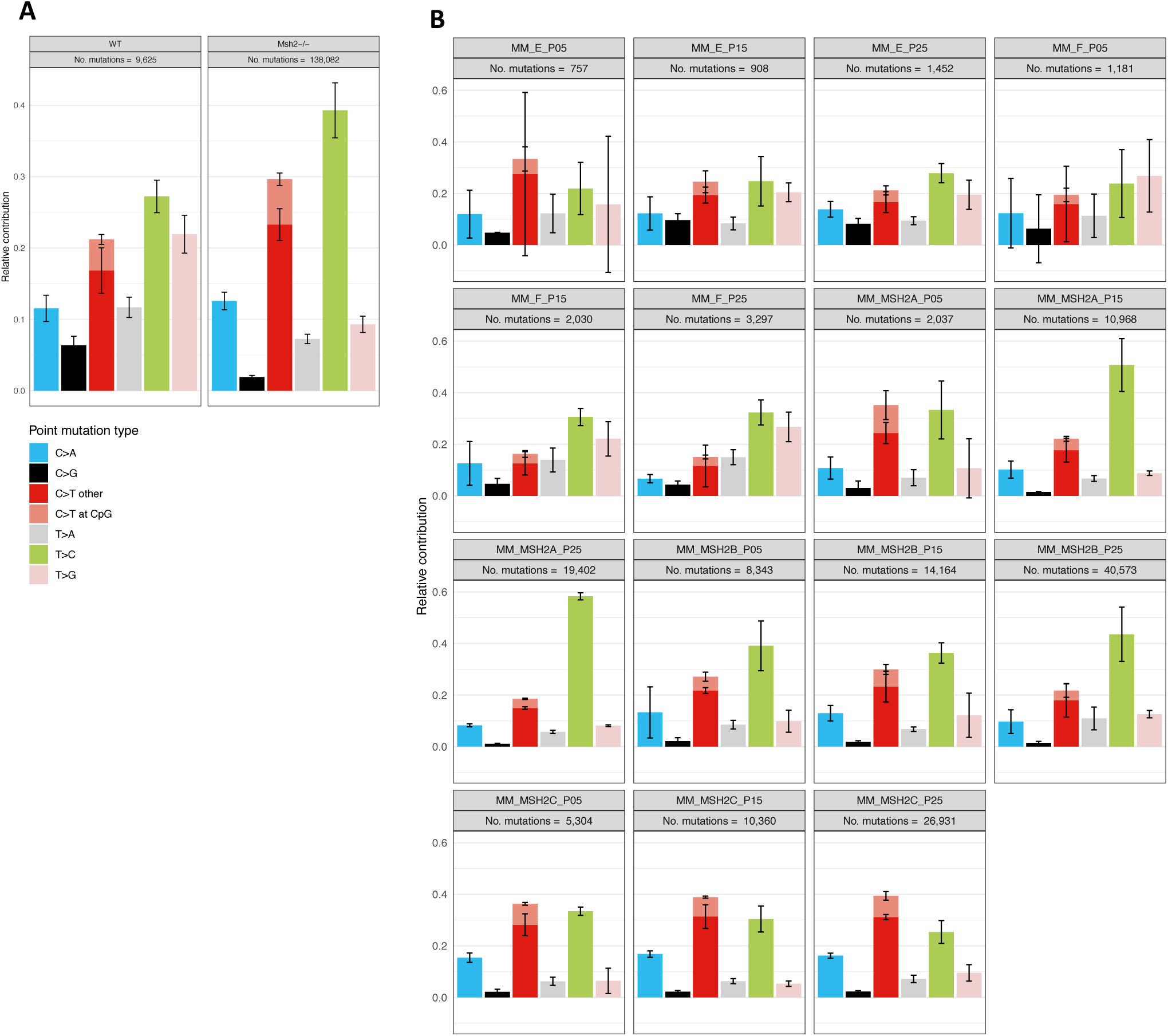
SNV spectra. (**A**) SNV spectra of cell strains of the two genotypes combined. (**B**) SNV spectra of each passage in each cell strain separately. Error bars present s.d.

**Figure S5.**
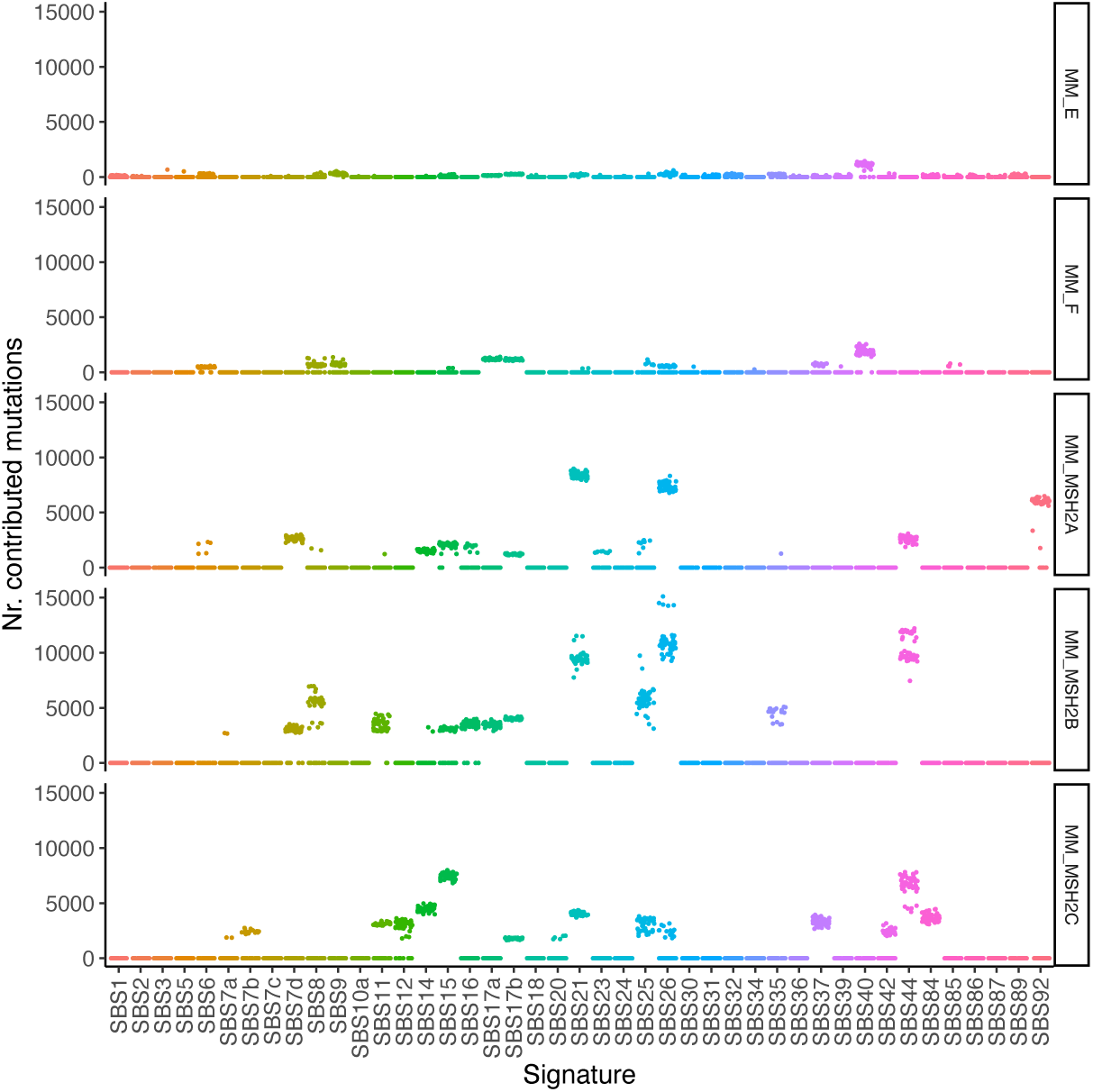
SNV signature refitted from COSMIC signatures. Each data point presents a cell. Only the COSMIC signatures estimated to contribute to at least one SNV were plotted. The analysis was performed using the “MutationalPatterns” package of R (Blokzijl *et al*., 2018).

**Figure S6.**
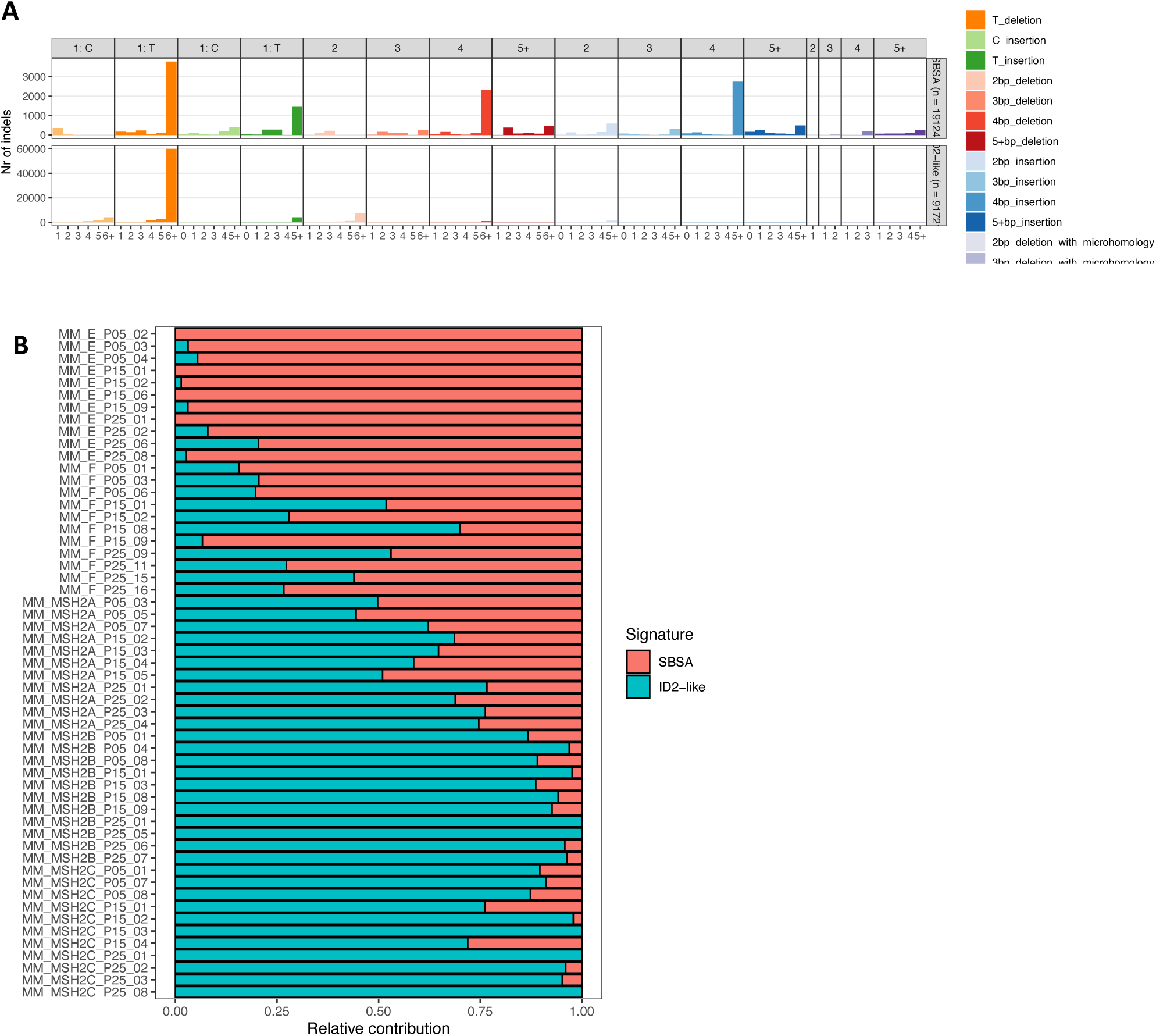
INDEL signatures. (**A**) Two INDEL signatures of the fibroblasts identified by *de novo* signature extraction using the “MutationalPatterns” package of R (Blokzijl *et al*., 2018). (**B**) Contribution of each INDEL signature to the total INDELs per cell.

**Figure S7.**
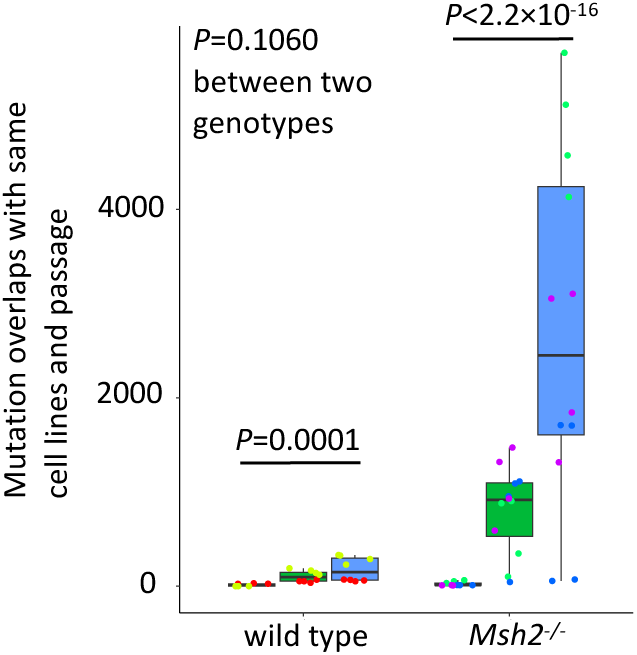
Mutational overlap per cell. Number of overlapping mutations among cells of the same passage and same cell strain (i.e., animal) to the number of overlapping mutations among cells of all cell strains. Each data point presents a cell. *P* values were estimated using linear mixed effects models, two-sided. Boxplot elements are defined as follows: center line indicates median, box limits indicate upper and lower quartiles, and whiskers indicate 1.5× interquartile range.

